# Regulation of Versican Expression in Macrophages is Mediated by Canonical Type I Interferon Signaling via ISGF3

**DOI:** 10.1101/2024.03.14.585097

**Authors:** Mary Y. Chang, Christina K. Chan, Jourdan E. Brune, Anne M. Manicone, Karol Bomsztyk, Charles W. Frevert, William A. Altemeier

**Author notes:** **Address correspondence to:** Mary Y. Chang, PhD, Dept. of Comparative Medicine, Campus Box 357340, University of Washington, Seattle, WA 98195-7340. M.Y.C and C.C. contributed equally to this work. **Additional contact information**: Christina K. Chan, Jourdan E. Brune, Anne Manicone, Karol Bomsztyk, Charles W. Frevert, DVM, ScD, William A. Altemeier, MD.

## Abstract

Growing evidence supports a role for versican as an important component of the inflammatory response, with both pro- and anti-inflammatory roles depending on the specific context of the system or disease under investigation. Our goal is to understand the regulation of macrophage-derived versican and the role it plays in innate immunity. In previous work, we showed that LPS triggers a signaling cascade involving TLR4, the Trif adaptor, type I interferons, and the type I interferon receptor, leading to increased versican expression by macrophages. In the present study, we used a combination of chromatin immunoprecipitation, siRNA, chemical inhibitors, and mouse model approaches to investigate the regulatory events downstream of the type I interferon receptor to better define the mechanism controlling versican expression. Results indicate that transcriptional regulation by canonical type I interferon signaling via the heterotrimeric transcription factor, ISGF3, controls versican expression in macrophages exposed to LPS. This pathway is not dependent on MAPK signaling, which has been shown to regulate versican expression in other cell types. The stability of versican mRNA may also contribute to prolonged versican expression in macrophages. These findings strongly support a role for macrophage-derived versican as a type I interferon-stimulated gene and further our understanding of versican’s role in regulating inflammation.

## Introduction

The extracellular matrix (ECM) has been shown to play critical roles in the inflammatory response.(Frevert et al., 2018; Gill et al., 2010; Kang et al., 2018; Sorokin, 2010; Vaday et al., 2001; Wight et al., 2020; Wight et al., 2014) The interactions between components of the ECM and leukocytes at sites of inflammation influences their adhesion, retention, migration and activation, with important consequences for disease progression and resolution. We are interested in the ECM component, versican, a chondroitin sulfate proteoglycan whose expression increases dramatically in a variety of acute and chronic pulmonary diseases, and in pulmonary cancers.(Andersson-Sjoland et al., 2015; Ayars et al., 2013; Bensadoun et al., 1996, 1997; Huang et al., 1999; Kim et al., 2009; Merrilees et al., 2008; Merrilees et al., 2004; Morales et al., 2011; Ricciardelli et al., 2009; Said et al., 2012; Weitoft et al., 2014) Versican is synthesized by multiple types of cells, including embryonic stem cells, epithelial, endothelial, stromal and neural cells, as well as by leukocytes.(Bode-Lesniewska et al., 1996; Brune et al., 2021; Chang et al., 2014; Hatano et al., 2012; Henderson & Copp, 1998; Kellar et al., 2021; LeBaron et al., 1992; Schonherr et al., 1991; Sotoodehnejadnematalahi et al., 2015) *In vitro* and *in vivo* studies have shown that this synthesis occurs in response to a variety of stimuli, including cytokines, growth factors, and agonists that bind to pattern recognition receptors. The roles of versican in these responses appear to be context-dependent, with both pro- and anti-inflammatory properties depending on the agonist, cellular involvement, kinetics of the immune response, and disease process.(Chang et al., 2017; Kang et al., 2018; Kang et al., 2017) Thus, elucidating the mechanisms that control versican expression in these specific contexts is essential for understanding this complex immunomodulatory molecule.

Several studies have established the foundation for understanding the regulatory mechanisms that control versican expression in normal primary cell cultures and in carcinoma-derived cell lines. The work of Schonherr, et. al., showed that genistein, a tyrosine kinase inhibitor, blocked the abilities of platelet-derived growth factor and transforming growth factor-beta to stimulate versican gene expression in arterial smooth muscle cells (SMCs).(Schonherr et al., 1991) The ability of angiotensin II (AngII) to stimulate versican expression in vascular SMCs was similarly shown to be inhibited by genistein. The effects of AngII on versican expression were also inhibited by an epidermal growth factor receptor (EGFR) tyrosine kinase inhibitor (AG1478) and a mitogen-activated protein kinase kinase (MAPK/MEK) inhibitor (PD98059), consistent with the known ability of AngII to stimulate transactivation of EGFR with downstream activation of the MAPK cascade.(Eguchi et al., 2001; Shimizu-Hirota et al., 2001) In addition, a phosphatidylinositol 3-kinase (PI3K) inhibitor (LY294002) was shown to block serum-induced versican promoter activation in aortic smooth muscle cells, and protein kinase B (PKB) was found to be an intermediary of this response.(Rahmani et al., 2005) These studies indicated that receptor tyrosine kinase-dependent signaling with downstream activation of PI3K and PKB is a common pathway for growth factor-mediated upregulation of versican expression in SMCs. A consequence of activation of PI3K/PKB signaling is the phosphorylation and inactivation of glycogen synthase kinase-3 beta (GSK-3β), which leads to stabilization and translocation of β-catenin to the nucleus where it forms a transcription factor complex with T-cell factor and lymphoid enhancer proteins (TCF/LEF) to activate target genes such as versican.(Maes et al., 2010; McCubrey et al., 2014)

Studies in carcinoma-derived cell lines also indicate a role for PKB signaling in the control of versican expression. In a colorectal carcinoma cell line, the tumor suppressor protein and transcription factor, p53, was shown to directly regulate versican transcription through a putative p53 consensus binding site in versican’s first intron.(Yoon et al., 2002) As with GSK-3β, p53 status depends on PKB signaling.(Brazil et al., 2002; Mayo & Donner, 2002) In other studies using teratocarcinoma cells as a model of embryonic stem cell development, microarray technology identified versican as one of fifty target genes upregulated early after stimulation with Wnt protein.(Willert et al., 2002) The Wnt signaling pathway converges with growth factor RTK signaling at the critical point of inactivation of GSK-3β, leading to subsequent β-catenin/TCF/LEF transcriptional activation of target genes.(Mayo & Donner, 2002; McCubrey et al., 2014; Nusse & Clevers, 2017) In another cancer model, SK-mel-131 melanoma cells, transcriptional regulation of versican is controlled by the c-jun N-terminal kinase (JNK) pathway, in which c-jun binds to an AP-1 site on the versican promoter.(Domenzain-Reyna et al., 2009) This response was inhibited not only by a JNK inhibitor (SP600125), but also by the MEK inhibitor (PD98059), confirming the unique intersection of JNK and MAPK/MEK pathways in melanoma cells. In addition, crosstalk between the MAPK/MEK pathway with β-catenin/TCF signaling was demonstrated in SK-mel-131 cells, providing further support for the important regulatory role of β-catenin in regulating versican expression.(Domenzain-Reyna et al., 2009)

In addition to transcriptional regulatory mechanisms, versican expression is influenced by post-transcriptional regulatory events. In *in vivo* studies of ovariectomized mice, estradiol was found to cause elongation of the poly(A) tail on versican, thereby increasing transcript stability and prolonging increased expression in uterine tissue.(Salgado et al., 2013) MicroRNAs can also control versican synthesis.(Rutnam et al., 2013) miR-143 attenuates versican expression and migration in smooth muscle cells, and miR-138 represses versican expression thereby aiding normal cardiac morphogenesis.(Morton et al., 2008; Wang et al., 2010)

We are interested in versican for its immunomodulatory properties in the context of acute pulmonary inflammation.(Brune et al., 2021; Chang et al., 2017; Chang et al., 2014; Kang et al., 2017; Kellar et al., 2021; Snyder et al., 2015) In our work, we have used *E. coli* lipopolysaccharide (LPS) and polyinosinic:polycytidilic acid (poly(I:C)) as molecular tools to dissect the signaling pathways regulating versican expression in primary bone marrow-derived macrophages from wild-type and genetically engineered mice.(Chang et al., 2017) Our findings show that versican expression in macrophages treated with LPS or poly(I:C) requires toll-like receptors (TLRs), the Trif adapter molecule, type I interferons (Ifn) and the type I Ifn receptor (Ifnar1), and support a role for macrophage-derived versican as an anti-inflammatory molecule. Interestingly, we found that an inhibitor of PI3K signaling (LY294002) caused a significant reduction in the versican response to LPS, but inhibitors of Wnt/β-catenin signaling (XAV939 or ICG001) had no impact on the ability of LPS to induce versican expression in macrophages.(Chang et al., 2017) We also observed that peak induction of versican in the lungs of mice infected with influenza A virus (IAV) correlated with peak induction of Ifn-β, and that versican expression was significantly attenuated in the lungs of IAV-infected mice lacking functional type I Ifn receptors (Ifnar1^-/-^) compared to IAV-infected wild-type mice.(Brune et al., 2021) Using dual *in situ* hybridization, versican expression was found to be decreased both in CD68^+^ myeloid cells and in PDGFRβ^+^ stromal cells in the lungs of IAV-infected Ifnar1^-/-^ versus wild-type mice. Taken together, these findings support a role for type I Ifn signaling in regulating versican expression in both myeloid and stromal cells.

The present study investigates the regulatory pathway downstream of type I Ifns and Ifnar1 to better define the mechanism controlling versican expression in macrophages. Using a combination of siRNA, chemical inhibitor, and mouse model approaches, we report that canonical type I Ifn signaling via interferon-stimulated gene factor 3 (ISGF3), the heterotrimeric transcription factor complex, controls the expression of versican in macrophages exposed to lipopolysaccharide (LPS). The detailed understanding of how versican expression is regulated in distinct cell types and models of inflammation may lead to unique approaches for targeting this immunomodulatory molecule’s pro- vs anti-inflammatory properties.

## Results

### The expression of versican RNA in bone marrow-derived macrophages in response to LPS is influenced by transcriptional and post-transcriptional mechanisms

As we have previously shown, versican is constitutively expressed at low levels in macrophages and is maximally induced at 8 hours after exposure to LPS (65.2 ± 6.0 –fold vs PBS control) (Figure 1A). We first used ChIP and qPCR to evaluate whether the increase in versican expression is due to transcriptional regulation by LPS. Analysis of RNA polymerase II (Pol II) binding at the Vcan gene (Prom 400 and Exon 1), estimated by antibodies to the C-terminal domain of Pol II, revealed increased recruitment of Pol II to the Vcan gene following exposure of macrophages to LPS (Prom400: 5.6-fold at 2 hours, 3.3-fold at 4 hours; Exon1: 4.4-fold at 2 hours and 5.5-fold at 4 hours) (Figure 1B). This indicates that transcription is occurring and is, at least in part, responsible for induction of versican expression in LPS-treated macrophages. Because versican mRNA continues to accumulate for up to 8-hours (Fig 1A), we evaluated whether transcription persisted throughout this duration. Macrophages were stimulated with LPS and then 4-hours later, exposed to actinomycin D, which binds to DNA and blocks the ability of RNA polymerase II to elongate nascent RNA molecules further. Following actinomycin D treatment, versican mRNA increased for an additional 1 hour and then remained stable for up to 8 hours. In contrast, versican continued to increase for 4-hours after vehicle treatment (8-hours after initial LPS stimulation) (Figure 1C). To verify the efficacy of actinomycin D, TATA binding protein (TBP) served as a control gene whose mRNA exhibited a classical decay curve following exposure to actinomycin D (Figure1D). These data indicate that transcription is an ongoing contributor to mRNA accumulation for 8 hours following LPS. These data also demonstrate that versican has a t_1/2_ > 8 hours indicating that versican mRNA has a slow decay rate. Because versican mRNA levels are very low in unstimulated macrophages, it is not possible to determine whether increase in mRNA stability contributes to mRNA accumulation after LPS exposure.

**Figure 1.**
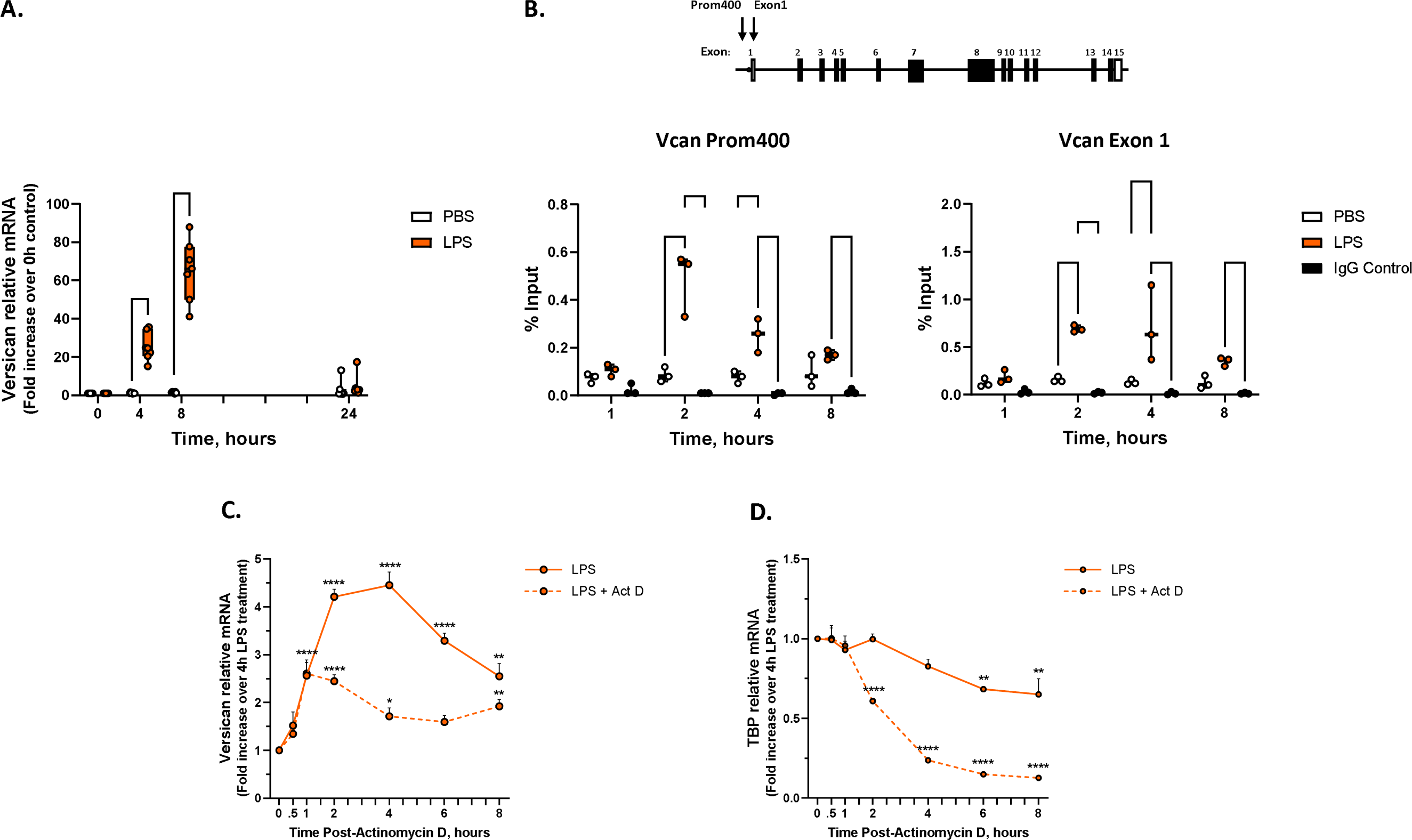
Evidence for transcriptional and post-transcriptional regulation of versican mRNA in macrophages. (A) BMDMs from wild-type mice were exposed to PBS or LPS (10 ng/ml) for up to 24 hours. Versican mRNA expression was evaluated by real-time qPCR. Data represent means ± SEM for cells from n = 7 mice. (B) Sheared cross-linked chromatin were assayed using an antibody to the C-terminal domain of RNA polymerase II. ChIP DNA was analyzed at the versican promoter (Prom 400) and first exon (Exon 1) by real-time qPCR. Data are expressed as % Input and represent means ± SEM n = 3 mice. A schematic of the versican gene (V0 variant) is shown: translated and untranslated exons are represented as solid and open rectangles, respectively; lines represent introns; arrows show locations of the amplicons. (C &D) BMDMs were treated with LPS for 4 hours and then exposed to vehicle (d^2^H_2_O) or actinomycin D (5 ug/ml). Cells were harvested at the indicated time points. QPCR data represent means ± SEM for n = 3 mice. Statistics were analyzed by 1-way ANOVA: * p < 0.05, ** p < 0.01, *** p< 0.001, **** p<0.0001.

### Versican induction directly depends on Ifn-β, but not Irf3 or Irf7

Our previous work also shows that Ifn-α and Ifn-β mimic the effects of LPS on versican expression by macrophages from wild-type (WT) mice and that versican mRNA is not increased in response to LPS in macrophages from mice lacking functional type I Ifn receptors (Ifnar1^-/-^), indicating that versican is regulated by type I Ifn signaling and is a type I Ifn-stimulated gene (ISG) in macrophages. As transcriptional activation of the Ifn-β gene requires the formation of an enhanceosome containing Irf3 and Irf7, we next used macrophages from mice deficient in both Irf3 and Irf7 (Irf3/7 dKO) to evaluate whether these transcription factors were essential for regulating versican expression.(Panne et al., 2007) For these experiments we compared the effects of 10 ng/ml of LPS to 100 U/ml of Ifn-β, which we have previously shown to induce comparable levels of versican in macrophages.(Chang et al., 2017) In WT macrophages, LPS causes a robust 1471.9 ± 306.2 –fold induction of Ifn-β mRNA, which peaks at 4 hours (Figure 2A) and precedes the peak induction of Vcan mRNA, which occurs at 8 hours (Figure 2B). Ifn-β causes a modest self-induction in a positive-feedback manner (Figure 2A) and a 56.5 ± 4.5 –fold induction of Vcan mRNA in WT macrophages (Figure 2B). The ability of LPS to induce Ifn-β is significantly attenuated in Irf3/7 dKO macrophages (∼97.5% reduction at 4 hours vs WT LPS); the self-induction of Ifn-β is similarly attenuated in Irf3/7 dKO macrophages (Figure 2A). Most notably, the ability of LPS to induce Vcan is significantly, but not totally, attenuated in Irf3/7 dKO macrophages (∼77.1% reduction at 8 hours vs WT LPS). Yet, Ifn-β induces Vcan mRNA to a similar degree in both WT and Irf3/7 dKO macrophages, in effect bypassing the lack of Irf3 and Irf7 (Figure 2B). These findings indicate that Irf3 and Irf7 do not directly activate versican expression in macrophages. However, Irf3 and Irf7 are necessary for the induction of Ifn-β, as has been well-described.(Honda et al., 2006) In short, LPS induction of Ifn-β is largely, though not entirely, responsible for increased versican expression in macrophages.LPS-mediated induction of versican is dependent on Jak1, but not Tyk2

**Figure 2.**
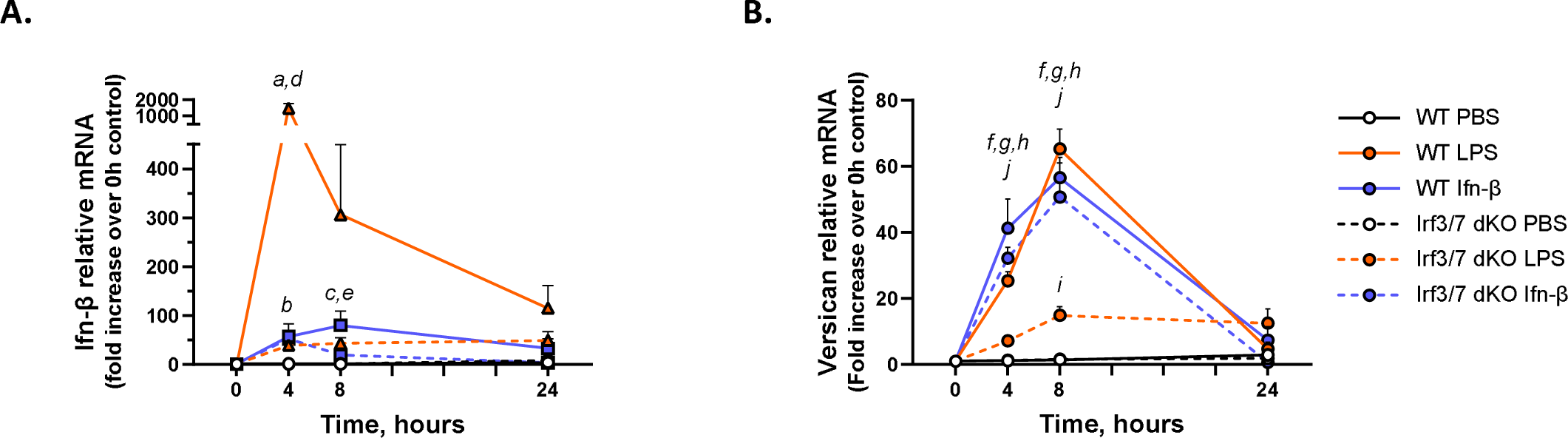
Roles of Ifn-b and Irf3/7 in the induction of versican mRNA. BMDMs from wild-type or Irf3/7 double knock-out mice were exposed to PBS, LPS (10 ng/ml), or Ifn-b (100 U/ml) for up to 24 hours. (A) Ifn-b and (B) versican mRNA expression were evaluated by real-time qPCR. Data represent means ± SEM for n = 5-7 mice. Statistics were analyzed by 2-way ANOVA with multiple comparisons: a, p < 0.0001 for WT LPS, 4h vs 0h control; b, p < 0.05 for WT Ifn-b, 4h vs 0h control; c, p < 0.001 for WT Ifn-b, 8h vs 0h control; d, p < 0.0001 for WT LPS vs Irf3/7 dKO LPS at 4h; e, p < 0.05 for WT Ifn-b vs Irf3/7 dKO Ifn-b at 8h; f, p < 0.0001 for WT LPS, 4 or 8h vs 0h control; g, p < 0.0001 for WT Ifn-b, 4 or 8h vs 0h control; h, p < 0.0001 for Irf3/7 dKO Ifn-b, 4 or 8h vs 0h control; i, p < 0.05 for Irf3/7 dKO LPS, 8h vs 0h control; j, p < 0.001 for WT LPS vs Irf3/7 dKO LPS at 4 or 8 h.

Type I Ifns mediate their biological activities by engaging cell surface Ifnar1 & 2 and activating downstream signaling events. Early events include receptor dimerization which induces phosphorylation and activation of their associated cytoplasmic Jak1 and Tyk2 enzymes. We used siRNA to investigate whether macrophage-derived versican expression depends on Jak1 and Tyk2 activation. Jak1 is constitutively expressed by macrophages, is not induced by either LPS or Ifn-β, and is effectively reduced at both the RNA (30nM siRNA: ∼69.0%; 300nM siRNA: ∼90.5% reduction vs control untransfected) and protein levels (300nM siRNA: ∼90%) by Jak1 siRNA (Figures 3A & 3B). Jak1 siRNA exerts a significant inhibitory effect on LPS-mediated induction of Ifn-β, and the self-induction of Ifn-β (Figure 3C). Similarly, Jak1 siRNA (300nM) effectively reduces the induction of versican in response to both LPS- and Ifn-β (100 U/ml), by 63.3 and 35.6%, respectively (Figure 3D). In separate experiments, we also showed that Jak1 siRNA attenuates the ability of a lower concentration of Ifn-β (10 U/ml) to induce versican expression (Supplemental Figure 1A). Tyk2 also is constitutively expressed by macrophages, but unlike Jak1, Tyk2 is induced by both LPS and Ifn-β (Figure 3E). Constitutive and inducible Tyk2 RNA and protein are inhibited by up to ∼80.1 and 60.0 %, respectively, with Tyk2 siRNA (Figures 3E & 3F). Tyk2 siRNA inhibits the self-induction of Ifn-β, but enhances LPS-induction of Ifn-β (2.0-fold at 30nM siRNA, p < 0.05) (Figure 3G). Tyk2 siRNA has no effect on LPS- or Ifn-β-mediated induction of versican (Figure 3H). To further evaluate the efficacy of Tyk2 silencing, we examined other genes described to be regulated by Tyk2 activity.(Poelzl et al., 2021) We verify here that LPS induces both IL1-β and IL18 in macrophages, and show that the increased expression of both cytokines is significantly reduced by Tyk2 siRNA (Figures 3I & 3J). These results indicate that LPS and Ifn-β both require Jak1 to induce versican expression in macrophages. Interestingly, the induction of versican by either agonist is not dependent on Tyk2. In addition, the stimulation of Ifn-β expression by LPS appears to be negatively regulated by Tyk2.

**Figure 3.**
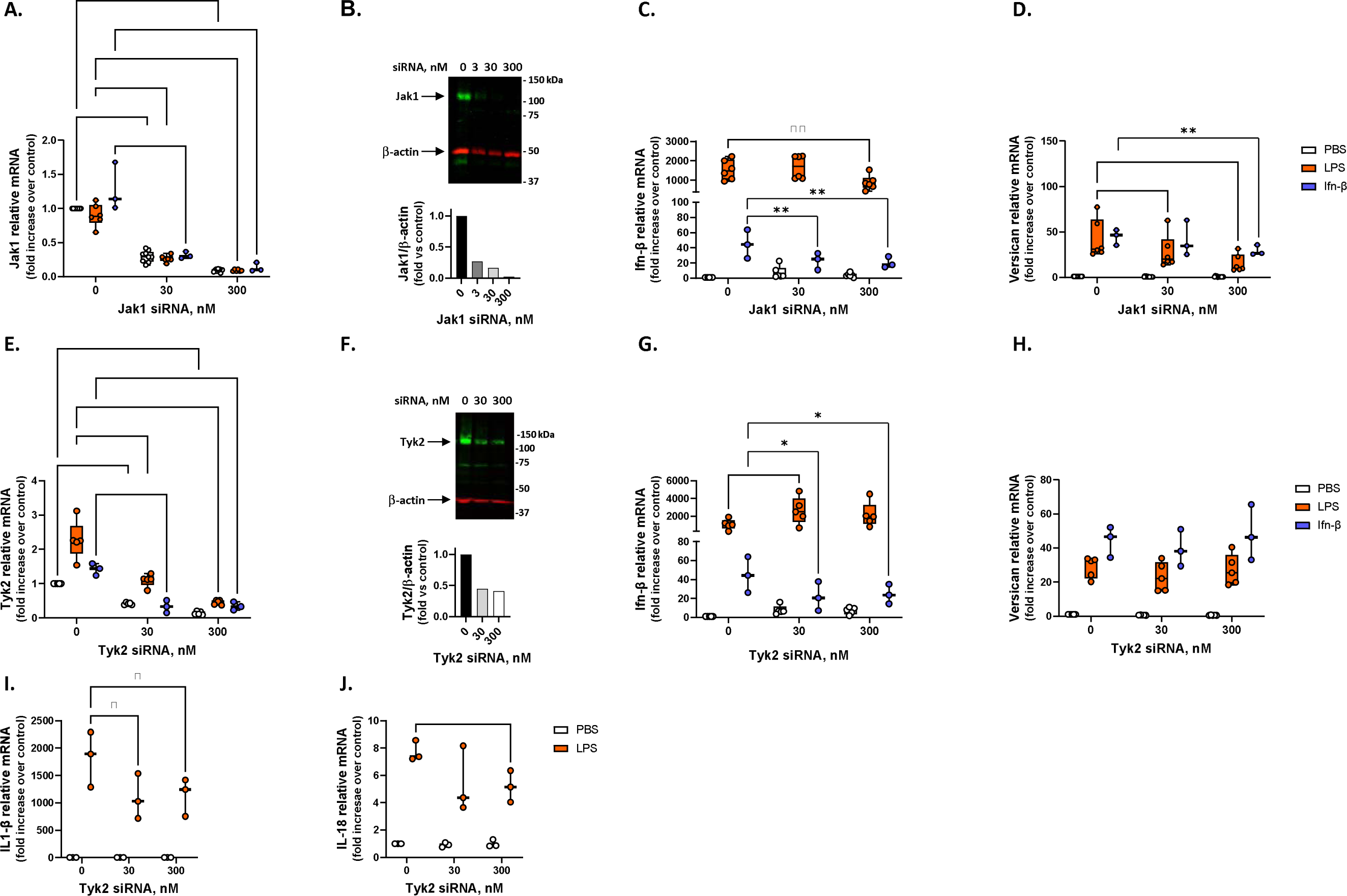
Effects of silencing Jak1 or Tyk2 on the induction of versican mRNA. BMDMs were transfected with 0, 30, or 300 nM of siRNA for (A-D) Jak1, or (E-J) Tyk2, for 24 hours prior to exposure to PBS, LPS (10 ng/ml), or Ifn-b (100 U/ml) for 4 hours. (A) Jak1, (E) Tyk2, (C & G) Ifn-b, (D & H) versican, (I) IL1-b and (J) IL18 mRNA expression were evaluated by real-time qPCR. Representative Western blots are shown for (B) Jak1 or (F) Tyk2 proteins with normalization to b-actin at 24 hours after transfection, with no stimulation. QPCR data represent means ± SEM for cells from n = 3-6 wild-type mice. Statistics were analyzed by 2-way ANOVA with multiple comparisons: * p < 0.05, ** p < 0.01, *** p< 0.001, **** p<0.0001.

### Silencing of Irf9 and Stat2, but not Stat1, inhibits LPS-mediated versican induction

In canonical type I Ifn signaling, the activation of Jak1 and Tyk2 leads to the recruitment of Stat1 and Stat2 monomers, which are phosphorylated and subsequently dimerize to facilitate binding of Irf9 in the cytoplasm, forming the interferon-stimulated gene factor 3 (ISGF3) complex.(Fu et al., 1990; Ivashkiv & Donlin, 2014; Muller et al., 1993; Owen et al., 2019) We also used siRNA to investigate whether macrophage-derived versican expression is dependent on ISGF3. Irf9, Stat1 and Stat2 RNA are all induced by LPS and Ifn-β in macrophages; these responses are inhibited by 50.2-72.2% following transfection with siRNA, as indicated (Figures 4A, 4E & 4I). siRNA also effectively inhibited constitutive protein expression (Figures 4B, 4F & 4J). While Irf9 siRNA inhibited the LPS-mediated induction of Ifn-β (Figure 4C), silencing of both Stat1 and Stat2 caused enhanced LPS induction of Ifn-β (Figure 4G & 4K). Irf9 siRNA inhibited both LPS- and Ifn-β-mediated induction of versican by 68.1 and 76.9%, respectively (Figure 4D). Stat2 siRNA also inhibited both LPS- and Ifn-β-mediated induction of versican by 23.5% and 28.8% at 300nM, respectively (Figure 4L). However, Stat1 siRNA had no effect on versican expression (Figure 4H). Similarly, siRNA for Irf9 and Stat2, but not Stat1, attenuated the ability of a lower concentration of Ifn-β (10 U/ml) to induce versican (Supplemental Figure 1B, 1C & 1D). This data indicates that the induction of versican by LPS or Ifn-β is dependent on Irf9 and Stat2. Furthermore, as observed with Tyk2, Stat1 and Stat2 also appear to negatively regulate Ifn-β expression.

**Figure 4.**
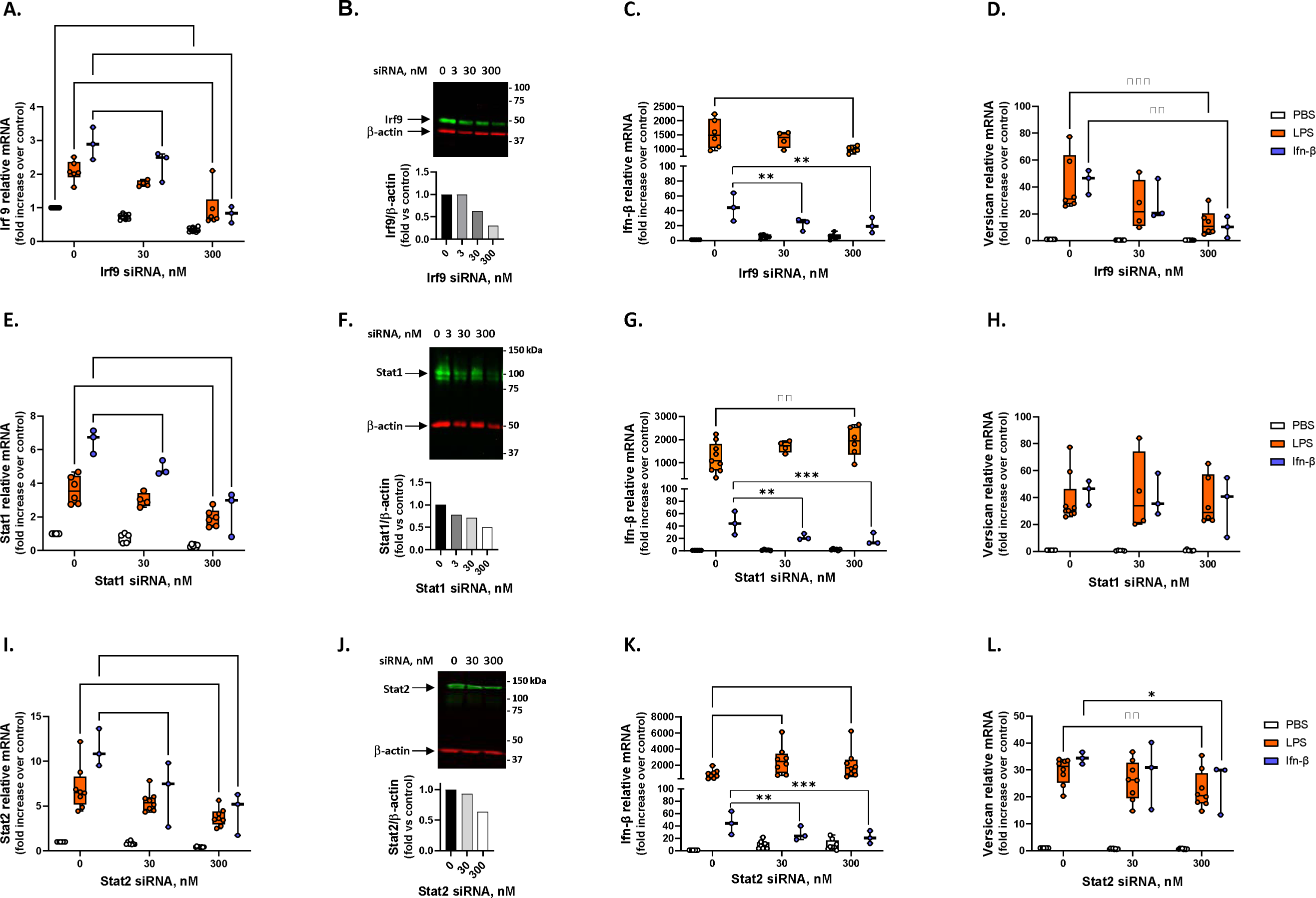
Effects of silencing Irf9, Stat1 or Stat2 on the induction of versican mRNA. BMDMs were transfected with 0, 30, or 300 nM of siRNA for (A-D) Irf9, (E-H) Stat1, or (I-L) Stat2 for 24 hours prior to exposure to PBS, LPS (10 ng/ml, or Ifn-b (100 U/ml) for 4 hours. (A) Irf9, (E) Stat1, (I) Stat2, (C, G & K) Ifn-β and (D, H & L) versican mRNA expression were evaluated by real-time qPCR. Representative Western blots are shown for (B) Irf9, (F) Stat1 or (J) Stat2 proteins with normalization to b-actin at 24 hours after transfection, with no stimulation. QPCR data represent means ± SEM for n = 3-9 wild-type mice. Statistics were analyzed by 2-way ANOVA with multiple comparisons: * p < 0.05, ** p < 0.01, *** p< 0.001, **** p<0.0001.

### Stat1 siRNA does not alter LPS-induced Stat1 phosphorylation

While transfection with Stat1 siRNA inhibited both constitutive and induced Stat1 mRNA as well as constitutive Stat1 protein, it did not inhibit the induction of versican in macrophages. We next examined the effect of siRNA on Stat1 phosphorylation, a necessary event for dimerization with Stat2 and subsequent interaction with Irf9. Western analysis showed that while Stat1 siRNA did cause a reduction in total Stat1 protein (normalized to β-actin) in LPS- or Ifn-β-treated macrophages, it had no effect on Stat1 phosphorylation (pStat1/Stat1) (Figure 5). These findings suggest that even with a reduction in Stat1 RNA and total protein, there was sufficient Stat1 phosphorylation to permit induction of versican in macrophages (Figure 4H).

**Figure 5.**
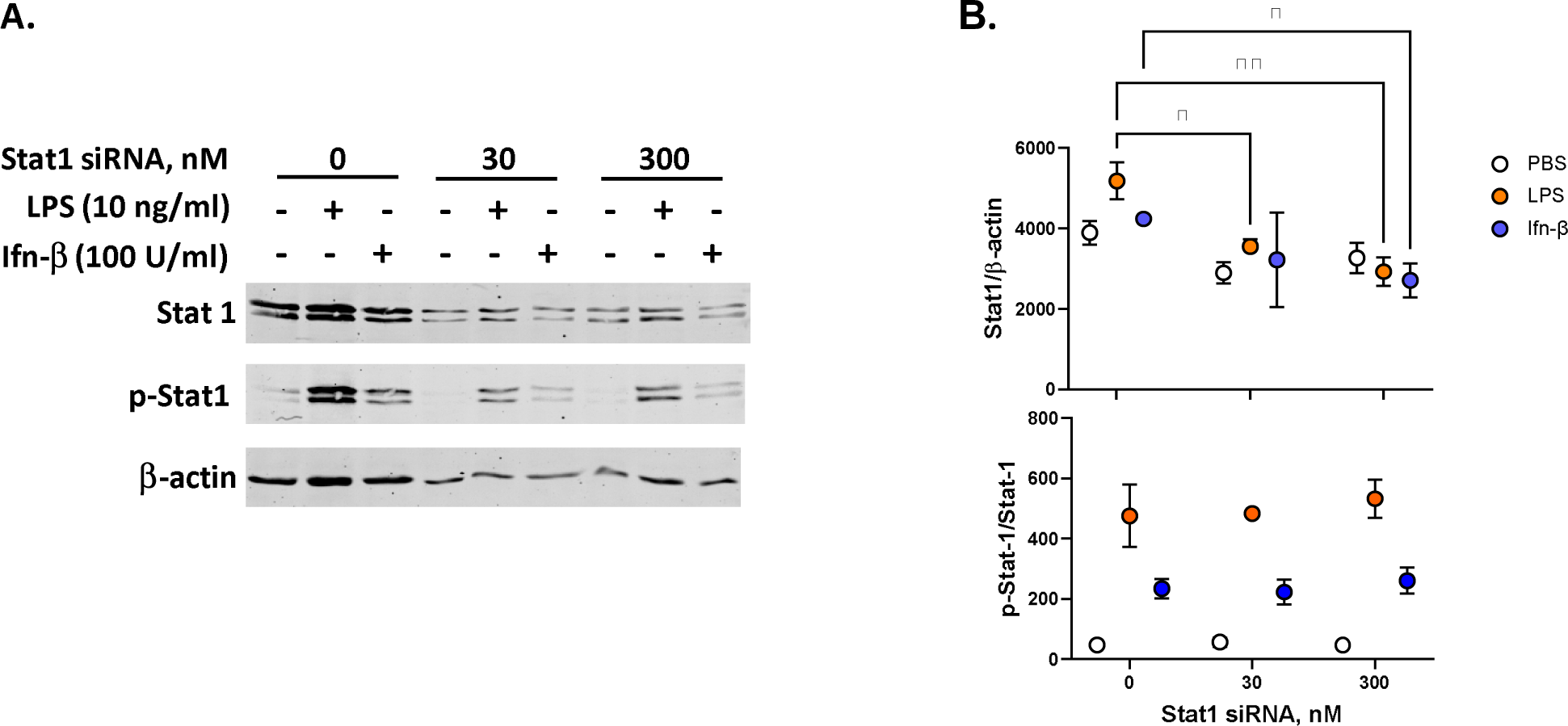
Effect of Stat1 siRNA on Stat 1 phosphorylation. BMDMs were transfected with 0, 30, or 300 nM of Stat1 siRNA for 24 hours prior to exposure to PBS, LPS (10 ng/ml), or Ifn-b (100 U/ml) for 4 hours. A. Representative Western blots are shown for Stat1, p-Stat1, and β-actin. B. Digital quantification data reflect means ± SD for cells from n = 2 wild-type mice. Statistics were analyzed by 2-way ANOVA with multiple comparisons: * p < 0.05, ** p < 0.01.

### Fludarabine inhibits LPS-induced Stat1 phosphorylation and versican expression

Fludarabine is a nucleoside analog that specifically inhibits the cytokine-induced activation of Stat1-dependent gene transcription; this effect has been attributed to the inhibition of Stat1 phosphorylation, both with and without concomitant inhibition of total Stat1 protein levels.(Frank et al., 1999) We used fludarabine to further examine the role of Stat1 in regulating versican expression. In macrophages, fludarabine slightly reduced the induction of Stat1 mRNA in response to 10 U/ml of Ifn-β but had no effect on the induction of Stat1 RNA (Figure 6A) or total protein (Stat1/β-actin) (Figure 6B) in response to LPS or 100 U/ml of Ifn-β. However, fludarabine caused an ∼27.6% reduction in Stat1 phosphorylation (pStat1/Stat1) (Figure 6B) and a comparable ∼26.5% reduction in versican RNA (Figure 6E), both in response to stimulation with LPS. Fludarabine also caused an ∼29.0% reduction in Stat1 phosphorylation induced by 10 U/ml of Ifn-β, though had no effect on pStat1/Stat1 in response to 100 U/ml of Ifn-β (Figure 6C). Even so, fludarabine inhibited versican induction in response to both 10 and 100 U/ml of Ifn-β by 40.5 and 20.9%, respectively, relative to PBS control-treated cells (Figure 6D). This data indicates that the induction of versican by LPS or Ifn-β depends on Stat1 phosphorylation. The inhibition of Stat1 phosphorylation caused enhanced LPS induction of Ifn-β (Figure 6C), as was also observed with silencing of Stat1 and Stat2 RNA (Figure 4G & 4H).

**Figure 6.**
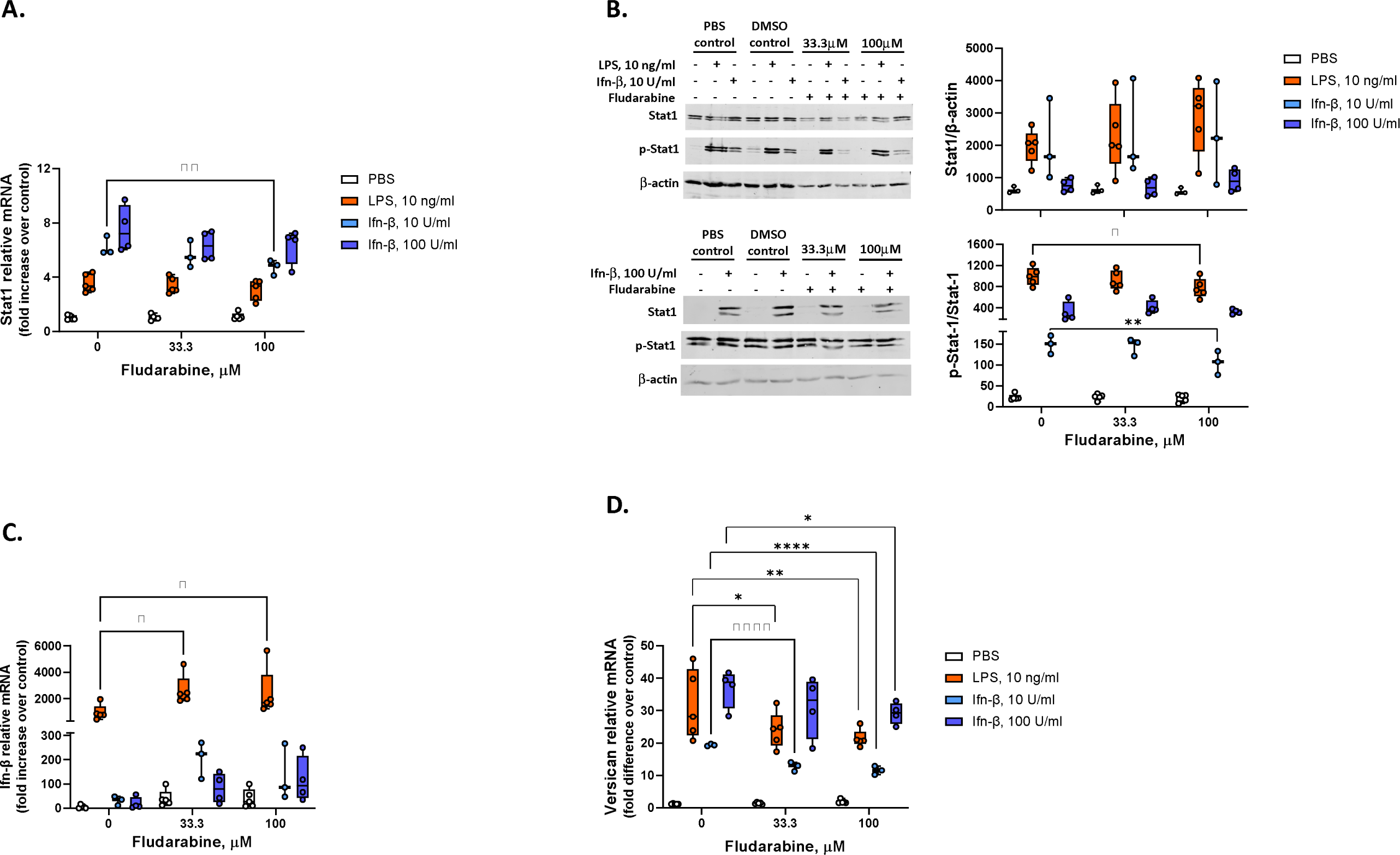
Role of Stat1 phosphorylation in the induction of versican mRNA. BMDMs were exposed to 0 (PBS or DMSO controls), 33.3 or 100 μM fludarabine for 24 hours prior to exposure to PBS, LPS, or Ifn-b (10 or 100 U/ml, as indicated) for 4 hours. (A) Stat1 mRNA expression was evaluated by real-time qPCR. (B,C) Western blot analyses were performed using antibodies to pStat1 (Tyr701), Stat1 and β-actin; representative blots are shown. Total Stat1 protein and the protein ratio of pStat1/total Stat1 were quantified by densitometric imaging. (D,E) Ifn-β and versican mRNA expression were evaluated by real-time qPCR. Data represent means ± SEM for cells from n = 3-6 wild-type mice. Statistics were analyzed by 2-way ANOVA with multiple comparisons versus the DMSO control (no differences between DMSO vs PBS): * p < 0.05, ** p < 0.01, *** p< 0.001, **** p<0.0001.

### Versican induction is not dependent on MEK1

Pharmacological inhibition of MEK1 and MEK2 inhibits versican expression in melanoma cells. (Domenzain-Reyna et al., 2009) Here, we investigated the role of MEK1 using bone marrow-derived macrophages from MEK1-deficient mice. First, we demonstrate that MEK1 protein is significantly decreased in macrophages from *Mek1^fl^LysM^Cre^* (MEK1-deficient) versus *Mek1^fl^* (control) mice (73.6% reduction) (Figure 7A). There were no differences in the abilities of LPS or Ifn-β to induce Ifn-β (Figure 7B) or versican (Figure 7C) expression in macrophages from *Mek1^fl^* versus *Mek1^fl^LysM^Cre^* mice. In contrast, the ability of LPS to stimulate Nos2 was reduced in *Mek1^fl^LysM^Cre^* cells (∼52.5% reduction vs *Mek1^fl^*), confirming that MEK1 deficiency decreases proinflammatory gene expression in bone marrow-derived macrophages (Figure 7D); this finding is similar to what has been described in alveolar macrophages.(Long et al., 2019) These results indicate that the induction of versican expression in macrophages treated with LPS or Ifn-β is not dependent on MEK1 signaling.

**Figure 7.**
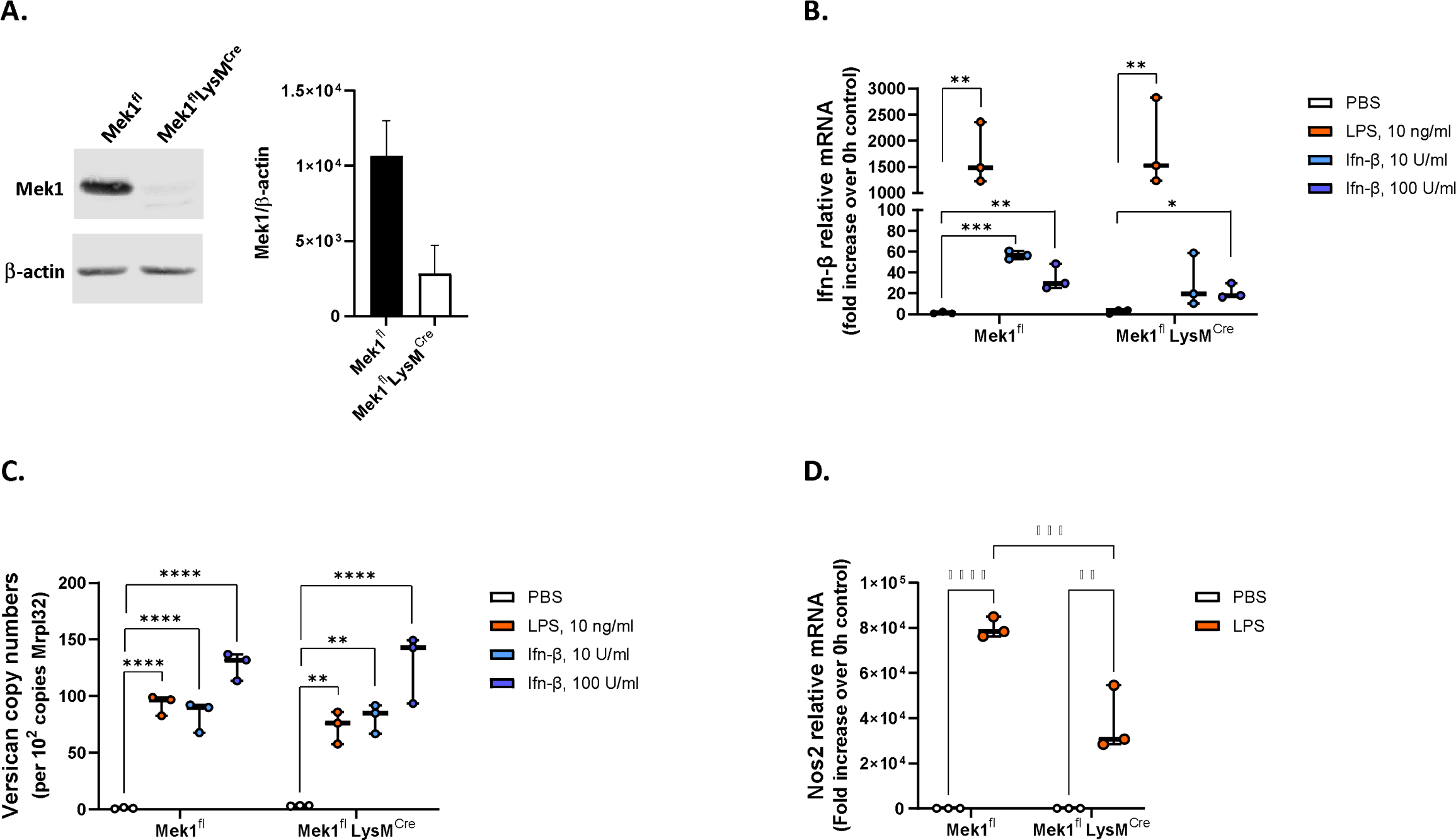
Role of Mek1 in the induction of Vcan. BMDMs from Mek1^fl^ or Mek1^fl^LysM^Cre^ mice were evaluated by Western blot analysis for Mek1 protein with normalization to b-actin. Representative blots with digital quantification of cells from replicate mice are shown (A). Cells were exposed to PBS, LPS, or Ifn-β for 4 hours and evaluated by real-time qPCR for (B) Ifn-β, (C) versican, or (D) Nos2 mRNA expression. Data represent means ± SEM for cells from n = 3 mice per strain. Statistics were analyzed by ANOVA with multiple comparisons: * p < 0.05, ** p < 0.01, **** p<0.0001.

## Discussion

We have previously reported that LPS triggers a signaling cascade involving TLR4, the Trif adaptor, type I Ifns, and Ifnar1, leading to the increased expression of versican by macrophages.(Chang et al., 2017) In the present study, we examined the signaling events downstream of Ifnar1 that regulate versican expression in more detail. We demonstrate the following findings. First, both transcriptional induction and transcript stability contribute to the level of versican expression in macrophages. Second, the transcription factors, Irf3 and 7, do not directly promote versican expression but are necessary for the induction of Ifn-β. Third, Ifn-β is essential for the induction of versican expression. Fourth, this response requires Jak1, Irf9, Stat1, and Stat2 proteins. These findings establish that LPS induces type I Ifn canonical signaling via the ISGF3 complex to regulate the expression of versican and establishes that macrophage-derived versican is a type I Ifn-stimulated gene (Figure 8).

**Figure 8.**
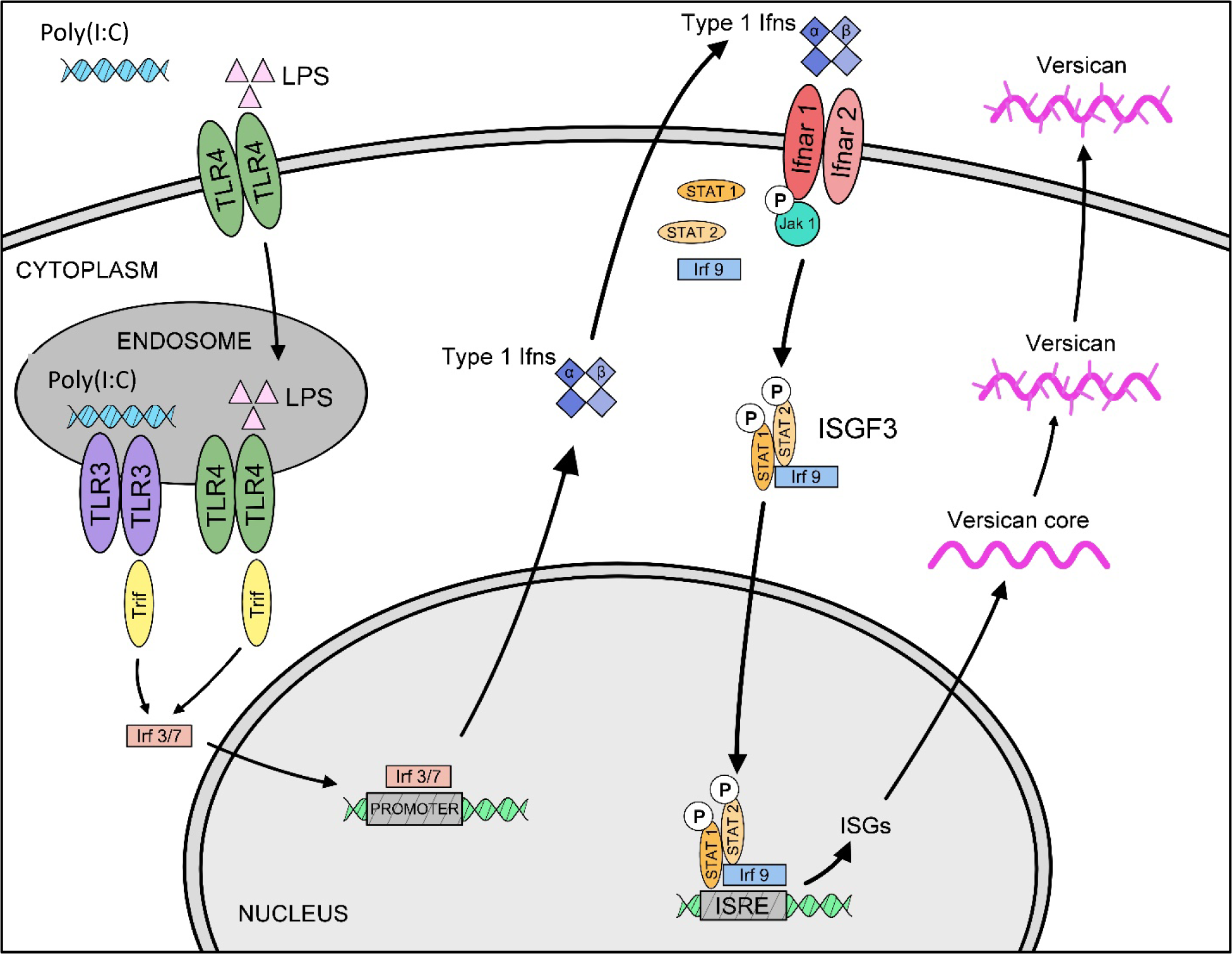
Schematic of regulatory events that control versican expression in macrophages. Previous work has shown that engagement of macrophage Toll-like receptors TLR4 and TLR3 by LPS and poly(I:C), respectively, result in enhanced versican expression. Subsequent to activation of TLR and TLR3, engagement of the TRIF adaptor molecule is known to activate transcription factors Irf3 and Irf7, leading to the production of Type I Ifns (Ifn-a/b) and recognition by type I Ifn receptors (Ifnar1/2).(Chang et al., 2017) We now identify Jak1, Irf9, Stat1 and Stat2 as essential signaling molecules downstream of Ifnar1/2 that mediate the induction of versican expression in macrophages.

Versican expression is regulated by transcriptional mechanisms in various primary smooth muscle cell cultures and carcinoma-derived cell lines.(Rahmani et al., 2005; Schonherr et al., 1991; Shimizu-Hirota et al., 2001; Willert et al., 2002; Yoon et al., 2002) These studies have shown that both growth factor and Wnt signaling pathways lead to the formation of the β-catenin/TCF/LEF transcription factor complex which binds to TCF/LEF binding motifs in the promoter region of the versican gene. Here we demonstrate that versican expression in macrophages is significantly influenced by transcriptional regulation (Figure 1). Additionally, versican mRNA is stable with a t_1/2_ > 8 hours, further contributing to mRNA accumulation after LPS. Building on our previous findings, we further establish that the transcriptional control of versican expression in macrophages is mediated by type I Ifn signaling and is distinct from the Wnt/β-catenin signaling pathway.

We evaluated whether specific transcription factors and signaling molecules are essential for the induction of versican expression in response to LPS compared to Ifn-β. The Irf transcription factors are considered master regulators of signaling by toll-like receptors (TLRs) and cytosolic pattern recognition receptors (PRRs). Of the nine Irf family members, Irf3 and Irf7 directly bind to promoter elements in type I Ifn genes in response to the engagement of TLR4 by bacterial LPS.(Honda & Taniguchi, 2006) Irf3 and Irf7 also bind to promoter elements in other target genes, many of which comprise the backbone of the innate immune response.(Hubel et al., 2019) Therefore, we used macrophages from Irf3/7 dKO mice to evaluate whether Irf3 and Irf7 directly regulate versican expression (Figure 2). We found that LPS did not induce Ifn-β expression in the absence of Irf3 and Irf7, as has been described.(Honda & Taniguchi, 2006; Sakaguchi et al., 2003). We also found that the ability of LPS to induce versican expression was largely, though not entirely, abrogated in Irf3/7 dKO macrophages. However, addition of exogenous Ifn-β could bypass the lack of Irf3/7 to induce versican expression. These findings indicate that the Irf3 and Irf7 transcription factors do not directly bind to the promoter region of versican to control gene expression. Rather, the regulation of versican expression is downstream of type I Ifn receptor signaling. Future studies will need to determine if the Irf3/7 independent pathway responsible for increased versican expression in macrophages treated with LPS is the result of non-canonical signaling through the type I Ifn receptor (Figure 2B).

The binding of type I Ifns to IfnarI and Ifnar2 initiates receptor dimerization, which results in the phosphorylation of their associated Janus kinases, Jak1 and Tyk2. This activates both canonical and non-canonical downstream signaling events. In the canonical pathway, three predominant complexes are formed that control distinct gene expression programs: Stat1, Stat2 and Irf9 form the heterotrimeric ISGF3 transcription factor complex which binds to Ifn-stimulated response elements (ISRE) sequences to activate classical anti-viral genes; Stat1 homodimers bind to gamma-activated sequences (GASs) to induce pro-inflammatory genes; and Stat3 homodimers indirectly suppress pro-inflammatory gene expression.(Ivashkiv & Donlin, 2014; Majoros et al., 2017; Owen et al., 2019) Here, we demonstrate that silencing of Jak1, Irf9 and Stat2, and interference with phosphorylation on Stat1, all inhibit the induction of versican expression in response to both Ifn-β and LPS (Figures 3-6). Therefore, we report the novel finding that macrophage-derived versican is induced by the ISGF3 arm of canonical type I Ifn signaling. One difference we find is that, unlike Jak1, Tyk2 is not essential for the LPS- or type I Ifn-mediated induction of versican in macrophages (Figure 3). While human cells fail to respond to type I Ifn in the absence of Tyk2, this is not the case for mouse studies.(Majoros et al., 2017) Signaling through type I Ifn receptors was found to be compromised but not abolished in Tyk2-deficient mice, with reduced levels of Stat1 protein but readily detectable activation of Stat1a and Stat1b in embryonic fibroblasts and bone marrow-derived macrophages treated with type I Ifns.(Karaghiosoff et al., 2000) The degree to which Tyk2 is necessary for mediating type I Ifn signaling may depend in part on the concentration used and the system under study, as low concentrations of Ifn-α (1-10 U/ml) had no anti-viral effect, while high concentrations (>100 U/ml) protected against the cytopathic effects of vesicular stomatitis virus on embryonic fibroblasts from Tyk2-deficient mice.(Shimoda et al., 2000) In previous work we compared the effects of low (10 U/ml) versus high (100 U/ml) concentrations of Ifn-β; we found that 100 U/ml induced a similar level of versican expression as did 10 ng/ml of LPS.(Chang et al., 2017) Therefore, for the majority of the present studies we employed 100 U/ml of Ifn-β. We did not investigate the effects of Tyk2 silencing in combination with a low concentration of Ifn-β. However, in a subset of experiments, we demonstrated that ISGF3 mediates the abilities of both low and high concentrations of Ifn-β to induce versican expression (Figure 6 and Supplemental Figure 1).

For most type I Ifn-stimulated genes, Jak-Stat signaling leading to the formation of the ISGF3 transcription factor complex is essential and sufficient for regulation. However, accessory signaling pathways are necessary for specific genes, and it has been proposed that the involvement of accessory pathways is cell type-specific.(Rani & Ransohoff, 2005) These pathways have been shown to involve PI3K, MAPK, p38 MAPK, nuclear factor-κB and IκB kinases.(David et al., 1995; Rani & Ransohoff, 2005) Of these, PI3K and MAPK are known to regulate versican expression in smooth muscle cells and in carcinoma cell lines. Both growth factor and Wnt signaling pathways require activation of PI3K for subsequent downstream formation of the β-catenin/TCF/LEF transcription factor complex, which binds to TCF/LEF binding motifs in the promoter regions of target genes, including versican.(Rahmani et al., 2005) Inhibition of the MAPK cascade using a pharmacological MEK inhibitor inhibited both epidermal growth factor receptor-mediated stimulation of versican expression and versican promoter activity in SK-mel-131 cells.(Domenzain-Reyna et al., 2009; Shimizu-Hirota et al., 2001) Taken together, these studies suggest that crosstalk between the MAPK and β-catenin/TCF/LEF pathways exerts a significant regulatory influence on versican expression. We have previously examined whether Wnt/β-catenin signaling might have an accessory role in the findings that we reported, and did not find that pharmacological inhibitors of Wnt signaling (XAV939 or ICG001) had an impact on the ability of LPS to induce versican expression in macrophages.(Chang et al., 2017) In the present study we used Mek1^fl^LysM^Cre^ mice to examine whether MAPK might have an accessory role, and show that MEK1 deficiency also has no impact on the ability of LPS to induce versican expression (Figure 7). These findings strongly suggest that regulation of versican by type I Ifn signaling and ISGF3 is a distinct pathway in macrophages.

Given that versican mRNA in macrophages continues to accumulate for 8 hours after LPS stimulation, we used an inhibitor of mRNA elongation, actinomycin D, four hours after LPS stimulation to determine whether this accumulation was dependent on transcription. Significant divergence in versican mRNA quantity between macrophages treated with actinomycin D and those treated with vehicle control indicates that transcription continues to contribute to versican mRNA accumulation for 8 hours after LPS exposure. We also report that versican mRNA is relatively stable with a half-life of greater than eight hours in macrophages (Figure 1). This finding is consistent with the general principles that: (i) mRNA species with long half-lives are enriched among genes related to metabolism and structure (e.g., extracellular matrix and cytoskeleton); and (ii) a major structural factor that affects mRNA stability is the presence of introns in genes (i.e., multi-exon genes), which makes mRNA more stable.(Sharova et al., 2009; Zhao & Hamilton, 2007) Versican is a large chondroitin sulfate proteoglycan with structural and regulatory functions.(Kang et al., 2018; Tang et al., 2022; Wight et al., 2020) It exists in four major isoforms (V0, V1, V2, and V3) generated by alternative splicing, with between 13-15 exons depending on the specific isoform. Thus, transcript stability contributes to the accumulation of versican in LPS-stimulated macrophages. However, as the constitutive level of versican is low in macrophages, we were not able to compare versican mRNA half-lives in untreated versus LPS-treated cells and cannot comment on whether LPS enhances versican mRNA stability. Future work will investigate potential mechanisms affecting versican mRNA stability, including poly(A) tail elongation, stabilizing RNA binding proteins, or microRNAs.

For most of the experiments, we compared the effects of 10 ng/ml of LPS to 100 U/ml of Ifn-β, which we had previously shown to induce comparable levels of versican in macrophages.(Chang et al., 2017) Our intent was to evaluate whether the effects of LPS on versican expression were mediated by Ifn-β. However, several interesting observations were made with respect to the regulation of Ifn-β expression. The inhibition of Stat1 phosphorylation caused enhanced LPS induction of Ifn-β (Figure 6), as was also observed with silencing of Stat1 and Stat2 RNA (Figure 4). These observations suggest that while Stat1 and Stat2 are required for the Ifn-β-dependent versican induction by LPS, they also appear to function as a negative regulatory feedback loop to limit Ifn-β expression. This could allow control of the amount or duration of gene expression for versican and other type I Ifn-stimulated genes following exposure to LPS. It will be of interest to examine this in more detail in future studies.

In previous work, we have shown not only that type I Ifns are necessary for versican expression in macrophages, but also that versican is necessary for type I Ifn production. We also showed that B6.LysMCre^+/+^/Vcan^fl/fl^ (LysM/Vcan^-/-^) mice, which are deficient in macrophage-derived versican, have increased recovery of inflammatory cells in the bronchoalveolar fluid (BALF) following treatment with poly(I:C), a TLR3/Trif agonist.(Chang et al., 2017) We now identify that the canonical type I Ifn pathway involving the ISGF3 transcription factor complex drives versican expression in macrophages. These findings suggest that macrophage-derived versican regulates inflammation and may play a role in limiting a potentially excessive inflammatory response. The finding that versican is necessary for type I Ifn production suggests a potential role for versican in viral control, though this was not examined in the present study.(Chang et al., 2017) Understanding whether this pathway is unique in macrophages is of great interest. Immunohistochemical findings from Brune, et. al., demonstrated that type I Ifn signaling increases versican expression and synthesis in lung stromal cells of C57BL6/J wild-type mice during influenza infection.(Brune et al., 2021) However, it is not known which arm of type I Ifn signaling is invoked in stromal cells in response to influenza – the formation of ISGF3 with anti-viral consequences or Stat1 homodimers which bind to gamma-activated sequences (GASs) to induce pro-inflammatory genes. Studies in B6.Rosa26-Cre^ERT+/-^Vcan^fl/fl^ (Vcan^-/-^) mice with a global deletion of versican, when treated with tamoxifen, showed decreased recovery of inflammatory cells in BALF following treatment with poly(I:C).(Kang et al., 2017) The contrasting results between LysM/Vcan^-/-^ versus Vcan^-/-^ mice exposed to poly(I:C) strongly suggest that versican’s role in the inflammatory response is contextual and could depend on its cellular source or the signaling pathway that is invoked. Adding to the complexity, versican function also depends on its degradation state, which affects its interactions with other extracellular matrix molecules.(Nandadasa et al., 2014) Extracellular matrix remodeling, driven in large part by the production of the ADAMTS-4 protease, has been shown to promote robust immune cell recruitment at the expense of lung function in response to severe influenza virus infection.(Boyd et al., 2020) This indicates a potentially significant role for versican degradation, a major substrate of ADAMTS-4, in the immunopathology of influenza viral infection.(Kelwick et al., 2015)

On-going studies in our laboratory using multiple models of cell-specific versican deficiency will aid in a better understanding of the contextual nature of versican’s role in the pulmonary innate immune response to influenza infection. A detailed understanding of the mechanisms regulating versican expression in different cells – macrophages versus fibroblasts - and inflammatory situations could lead to unique approaches for modifying versican’s anti- or pro-inflammatory properties.

## Methods

### Reagents

LPS from E. coli serotype 0111:B4 was purchased from List Biological Laboratories (Campbell, CA). Ifn-b was from PBL Interferon Source (Piscataway, NJ). FlexiTube GeneSolution siRNAs for JAK1 (GS16451), TYK2 (GS54721), Irf9 (GS16391), STAT1 (GS20846), and STAT2 (GS20847) were from Qiagen (Germantown, MD). Fludarabine was from Selleckchem (Houston, TX). β-mercaptoethanol and actinomycin D were from Sigma (Burlington, MA). RIPA buffer, Halt proteases with phosphatase inhibitors and high-capacity cDNA archive kits were from ThermoFisher Scientific (Grand Island, NY). AquaBlock was from Arlington Scientific (Springville, UT). The RNeasy Plus Mini Kit was from Qiagen (Germantown, MD). The ChIP-IT PBMC Kit was from Active Motif (Carlsbad, CA). The Amaxa Mouse Macrophage Nucleofector Kit was from Lonza (Cologne, Germany). Gene-specific TaqMan primer-probe mixes used for quantitative real-time polymerase chain reaction (PCR) of versican (Assay ID Mm01283063_m1), Ifn-b (Mm00439552_s1), Jak1 (Mm00600614_m1), Tyk2 (Mm00444469_m1), Irf9 (Mm00492679_m1), Stat1 (Mm01257286_m1), Stat2 (Mm00490880_m1), IL1-b (Mm00434225_m1), IL18 (Mm00434225_m1), NOS 2 (Mm00440502_m1), TBP (Mm01277042_m1), and Mrpl32 (Mm00777741_sH) were from ThermoFisher Scientific (Grand Island, NY). Antibodies to RNA polymerase II (14958S), Jak1 (3332S), Tyk2 (35615S), Irf9 (28845S), Stat1 (9172S), pStat1(9167S), Stat2 (72604S), and MEK1 (12671S) were from Cell Signaling Technology (Danvers, MA). Antibodies to b-actin (ab8226) were from Abcam (Waltham, MA). The rabbit IgG control (I-1000-5) was from Vector Laboratories (Newark, CA).

### Mice

Wild-type C57BL/6 were purchased from Jackson Laboratories (Bar Harbor, ME). Irf3/7 double knock-out (dKO) mice were a generous gift of Dr. Michael Gale (Seattle, WA). Mek1^fl^ and Mek1^fl^LysM^Cre^ mice on a C57BL/6 background were a generous gift of Dr. Anne Manicone (Seattle, WA). All mice were housed in a specific pathogen-free animal facility until the day of the experiment. The room temperature was held in a range of 68-79°F, with the goal of 72°F and an acceptable temperature variation of no more than 4°F over a 24 hour period. The acceptable room humidity range was 30-70%. Room lighting was programmed for 14 hours of light and 10 hours of dark. Mice were housed in HEPA-filtered ventilated shoe box cages (Allentown, PA), and the ventilation in the animal housing room was at least 10-15 fresh air changes per hour per room. All mice were provided environmental enrichment, fed rodent diet ad libitum, and had free access to fresh water at all times. Daily health checks were performed by the UW husbandry staff, who are overseen by laboratory animal veterinarians. The Institutional Animal Care and Use Committee (IACUC) of the University of Washington (UW) approved all experiments.

### Cell Culture Studies

Bone marrow was collected from femurs and tibia of C57BL/6, Irf3/7 dKO, Mek1^fl^, and Mek1^fl^LysM^Cre^ mice. Bone marrow cells were then cultured in “macrophage medium” (RPMI 1640, 10% FCS, 30% L929 cell supernatant, 2 mM L-glutamine, 100 IU/ml penicillin, and 100 g/ml streptomycin) for 6 days.(Tanino et al., 2012) For qPCR studies, macrophages were re-plated in macrophage medium in 12-well tissue culture dishes at a density of 0.44 x 10^6^ cells/well for 24 hours before stimulation in RPMI containing 10% FBS for up to 48 hours. For ChIP studies, macrophages were re-plated in macrophage medium in 100mm dishes at a density of 8 x 10^6^ cells/well for 24 hours.

### Chromatin Preparation and ChIP Assay

Macrophages were stimulated in the presence or absence of LPS (10 ng/mL) in RPMI containing 10% FBS for up to 8 hours. Media were removed to a fresh tube, cells were incubated with PBS on ice for 30 minutes, and cells were harvested. Media and cell layer were pooled and centrifuged at 400 x g for 10 minutes. After aspiration, cells were cross-linked in 1% formaldehyde/PBS for 10 minutes, quenched using a final concentration of 125 mM glycine for 5 minutes and washed twice with PBS. Cross-linked cells were then lysed using multiple incubations with cell membrane lysis buffer (5mM PIPES pH8.0, 85mM KCl, 0.5% NP-40, 0.5% TX-100) and nuclear membrane lysis buffer (PBS pH7.4, 1% NP-40, 0.5% sodium deoxycholate, 0.1% SDS).(Browne et al., 2014) After lysis, cells were pelleted by centrifugation (850 x g for 5 minutes at 4^0^C), the supernatant was removed and the cells were re-suspended in chromatin shearing buffer.(Bomsztyk et al., 2019) The chromatin was subjected to 3 x 10 minute cycles of ultrasonic shearing using a PIXUL multi-sample sonicator (Matchstick Technologies Inc., Kirkland, WA, and Active Motif, Carlsbad, CA).(Bomsztyk et al., 2019) With this procedure, chromatin was sheared to fragments of 500-1,000 base pairs in length, as confirmed using agarose gel electrophoresis. Soluble chromatin was immunoprecipitated using specific antibodies to RNA Polymerase II and a rabbit IgG control. Chromatin-antibody complexes were purified using ChIP-IT Protein G Magnetic beads and a high/low salt buffer (20 mM Tris-HCl, pH8.0, 2 mM EDTA, 1% NP-40, 0.1% SDS, 0.5 mM /0.15 M NaCl). Chromatin was eluted with 0.1N NaHCO3 and 1% SDS followed by RNase and Proteinase K treatment. DNA was purified using a QIAquick PCR Purification kit and qPCR was performed with Power Up SyBr Green PCR master mix using the ABI 7900HT Real Time PCR system (Applied Biosystems, Waltham, MA). PCR reactions contained primers for murine Vcan Prom400 (mVCAN Prom 400: forward, 5’-AGT ATC TCT TTC AGG TTG GCA -3’; reverse, 5’-TGT TTA ATG CGA CCA CGG AG -3’) or Exon 1(mVCAN Ex1: forward, 5’-CAC AAC CCG CAT TTG AAC TT -3’; reverse, 5’-CAC AGC ACC TAA TGC TCA CA -3’) at a 300 nM concentration and 5 ul of IP sample/reaction. PCR conditions were set as follows: 50°C for 2 minutes, 95°C for 2 minutes (denature at 95°C for 15 seconds, primer anneal at 55°C for 15 seconds, 72°C for 60 seconds), with 40 cycles. Data was calculated using the percent input method, in which the amount of target sequence measured in the immunoprecipitate isolate is compared to the total amount of the target sequence in the input isolate using the following equation: % Input = 2^((Ct(IN)-Log^_2_^(DF))-Ct(IP))^ * 100, where Ct(IN) is the threshold cycle value for the input sample, Ct(IP) is the threshold cycle value for the immunoprecipitate sample, and DF is the dilution factor which was 50 when 2% of the starting chromatin was used for the input sample.

### Transfections with siRNA

BMDMs were transfected with siRNA using the Lonza Mouse Macrophage Nucleofector kit, per the manufacturer’s instructions. In brief, adherent macrophages were placed on ice for 30 minutes and then were gently removed from tissue culture dishes using cell lifters (source). 1 x 10^6^ cells per sample were transfected with 0-300 μM of siRNA in 100 μl of Nucleofector Solution using the Y-001 Nucleofector Program on the Lonza Nucleofector II device. Cells were transferred to 12-well tissue culture trays and incubated at 37°C with 5% CO_2_ for 24 hours.

### Inhibition of transcription

Actinomycin D was dissolved slowly in d^2^H_2_O for a 2 mg/ml stock solution and stored at 4°C.(Lai et al., 2019) Macrophages were stimulated with PBS, LPS (10 ng/mL), or Ifn-β (100 U/ml) for 4 hours, after which actinomycin D (final concentration, 5 ug/ml) or vehicle (d^2^H_2_O) were added. Cells were harvested at 0, 0.5, 1, 2, 4, 6 and 8 hours post- actinomycin D for the isolation of total RNA and quantitative real-time PCR. The average threshold cycle (Ct) at each time point was normalized to the average Ct at 4 hours of stimulation with LPS or Ifn-b to obtain the ΔCt value. Relative abundance was then calculated as 2^(-ΔCt)^.(Ratnadiwakara & Anko, 2018)

### Isolation of RNA and quantitative real-time PCR

Total RNA was obtained from cell culture monolayers with RNeasy Plus Mini Kits. RNA was reverse transcribed with the High-Capacity cDNA Archive Kit at 25°C for 10 min, at 37°C for 2 h, and at 90°C for 5 min. The resulting cDNA was used for quantitative real-time PCR. Real-time PCR was carried out in a total volume of 25 μl with a master mixture including all reagents required for PCR and gene-specific TaqMan primer-probe mixes. Cycle parameters were 50°C for 2 minutes, 95°C for 10 minutes, followed by 40 cycles of 95°C for 15 seconds and 60°C for 1 minute; the ABI PRISM 7000 Sequence Detector (Applied Biosystems) was used. The threshold cycle (Ct) was calculated as the difference in Ct for the target genes versus Mrpl32, and the relative mRNA expression was expressed as fold increase over the values obtained from RNA collected from control cells.

### Western immunoblotting

Cells were lysed in Pierce RIPA buffer containing Halt protease and Phosphatase Inhibitor and 1 mM PMSF. Proteins (30ug) were separated on either 8% SDS-PAGE gels or 10% SDS-Page gel. After semi-dry transfer of protein gels to nitrocellulose, membranes were blocked with 10% AquaBlock in Tris-buffered saline containing 0.02% Tween 20 (TBST) and incubated with primary antibodies specific for STAT1 (1:1000), Phospho-STAT1(tyrosine 701) (1:1000), JAK1 (1:1000), STAT 2(1:1000), Irf 9 (1:1000), and β-actin (1:2000) at 4 °C overnight. Membranes were washed four times in TBST and incubated with fluorescently-labeled secondary antibodies (1:20,000; LI-COR Biotechnology, Lincoln, NE) for 1 h. Membranes were washed again and scanned in an Odyssey M imaging system (LI-COR Biotechnology, Lincoln, NE). Densitometric analysis of proteins bands was carried out using the Empiria studio software provided by LI-COR. Cellular protein loading was normalized to β-actin.

### Statistical analysis

Values are reported as Means ± SEM. Significance was evaluated by one- or two-way analysis of variance with multiple comparisons, as appropriate.

## Acknowledgements

The authors would like to thank Daniel Mar (Research Scientist, University of Washington) for assistance with ChIP analyses and Peter Waldron (Research Scientist, University of Washington) for assistance with animal care and procedures.

