## Supplemental Figure 1 for "Regulation of Versican Expression in Macrophages is Mediated by Canonical Type I Interferon Signaling via ISGF3"

### Slide 1
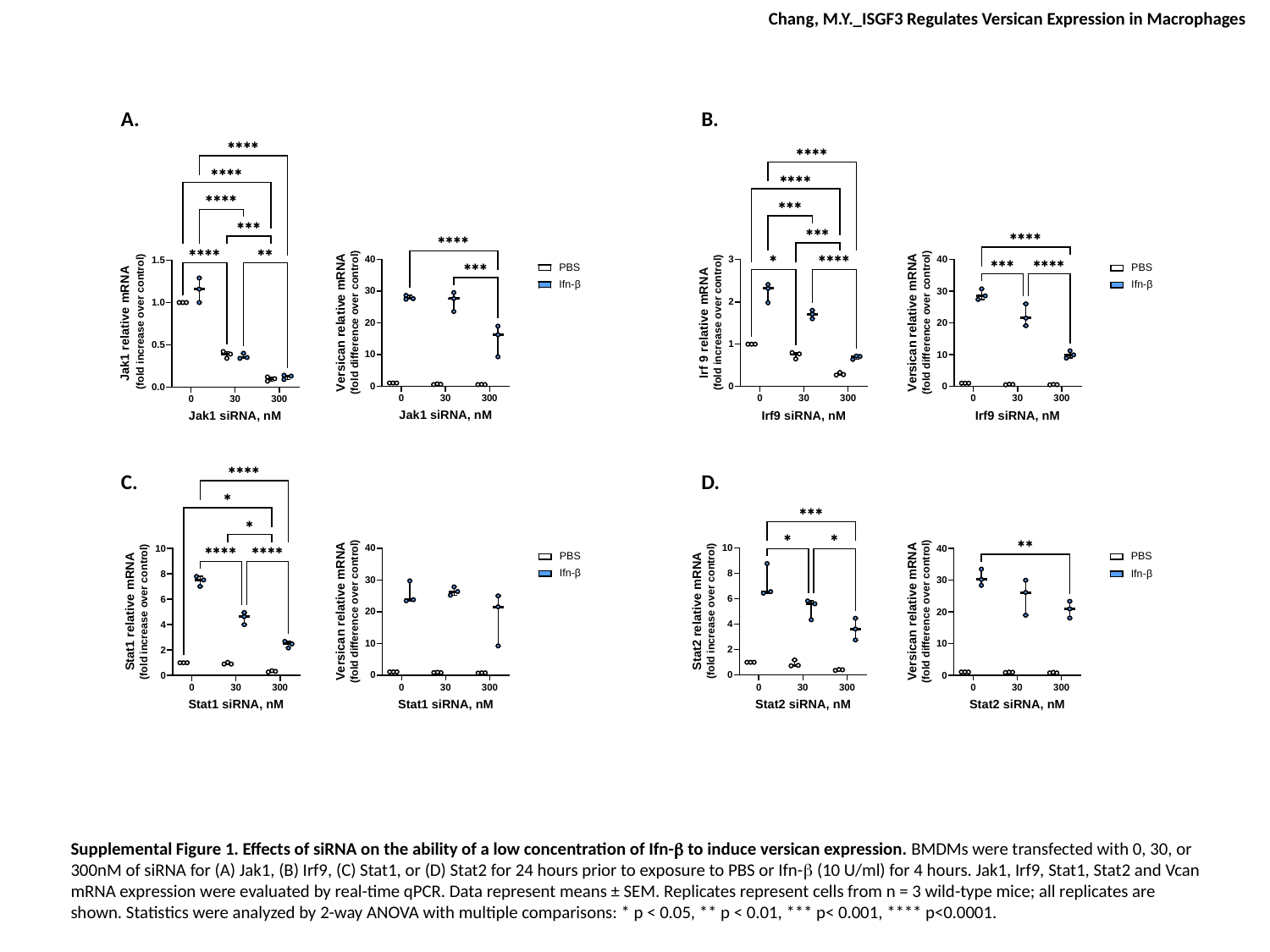

Chang, M.Y._ISGF3 Regulates Versican Expression in Macrophages
A.
B.
C.
D.
Supplemental Figure 1. Effects of siRNA on the ability of a low concentration of Ifn-β to induce versican expression. BMDMs were transfected with 0, 30, or 300nM of siRNA for (A) Jak1, (B) Irf9, (C) Stat1, or (D) Stat2 for 24 hours prior to exposure to PBS or Ifn-β (10 U/ml) for 4 hours. Jak1, Irf9, Stat1, Stat2 and Vcan mRNA expression were evaluated by real-time qPCR. Data represent means ± SEM. Replicates represent cells from n = 3 wild-type mice; all replicates are shown. Statistics were analyzed by 2-way ANOVA with multiple comparisons: * p < 0.05, ** p < 0.01, *** p< 0.001, **** p<0.0001.
